# The Distal Appendage Protein CEP164 Is Essential for Efferent Duct Multiciliogenesis and Male Fertility

**DOI:** 10.1101/2020.11.26.399642

**Authors:** Mohammed Hoque, Danny Chen, Rex A. Hess, Feng-Qian Li, Ken-Ichi Takemaru

## Abstract

Cilia are evolutionarily conserved microtubule-based structures that perform diverse biological functions. Cilia are assembled on basal bodies and anchored to the plasma membrane via distal appendages. Multiciliated cells (MCCs) are a specialized cell type with hundreds of motile multicilia, lining the brain ventricles, airways, and reproductive tracts to propel fluids/substances across the epithelial surface. In the male reproductive tract, MCCs in efferent ducts (EDs) move in a whip-like motion to stir the luminal contents and prevent sperm agglutination. Previously, we demonstrated that the essential distal appendage protein CEP164 recruits Chibby1 (Cby1), a small coiled-coil-containing protein, to basal bodies to facilitate basal body docking and ciliogenesis. Mice lacking CEP164 in MCCs (FoxJ1-Cre;CEP164^fl/fl^) show a significant loss of multicilia in the trachea, oviduct, and ependyma. In addition, we observed male sterility, however, the precise role of CEP164 in male fertility remained unknown. Here, we report that the seminiferous tubules and rete testis of FoxJ1-Cre;CEP164^fl/fl^ mice exhibit substantial dilation, indicative of dysfunctional multicilia in the EDs. Consistent with these findings, multicilia were hardly detectable in the EDs of FoxJ1-Cre;CEP164^fl/fl^ mice although FoxJ1-positive immature cells were present. Sperm aggregation and agglutination were commonly noticeable in the lumen of the seminiferous tubules and EDs of FoxJ1-Cre;CEP164^fl/fl^ mice. In FoxJ1-Cre;CEP164^fl/fl^ mice, the apical localization of Cby1 and the transition zone marker NPHP1 was severely diminished, suggesting basal body docking defects. TEM analysis of EDs further confirmed basal body accumulation in the cytoplasm of MCCs. Collectively, we conclude that deletion of CEP164 in the MCCs of EDs causes basal body docking defects and loss of multicilia, leading to sperm agglutination, obstruction of EDs, and male infertility. Our study therefore unravels an essential role of the distal appendage protein CEP164 in male fertility.

**Author Summary:** Multicilia are tinny hair-like microtubule-based structures that beat in a whip-like pattern to generate a fluid flow on the apical cell surface. Multiciliated cells are essential for the proper function of major organs such as brain, airway, and reproductive tracts. In the male reproductive system, multiciliated cells are present in the efferent ducts, which are small tubules that connect the testis to the epididymis. However, the importance of multiciliated cells in male fertility remains poorly understood. Here, we investigated the role of the critical ciliary protein CEP164 in male fertility using a mouse model lacking CEP164 in multiciliated cells. Male mice are infertile with reduced sperm counts. We demonstrate that, in the absence of CEP164, multiciliated cells are present in the efferent ducts but fail to extend multicilia due to basal body docking defects. Consistent with this, the recruitment of key ciliary proteins is perturbed. As a result, these mice show sperm agglutination, obstruction of sperm transport, and degeneration of germ cells in the testis, leading to infertility. Our study therefore reveals essential roles of CEP164 in the formation of multicilia in the efferent ducts and male fertility.

## Introduction

Spermatogenesis is a highly complex and finely tuned process in the testis that begins with a diploid stem cell and ends with four haploid germ cells after two successive meiotic divisions [1]. Spermatogenesis can be divided into three broad steps: (1) an initial asymmetric mitotic division, where spermatogonia give rise to primary spermatocytes, (2) two meiotic cell divisions, where these primary spermatocytes produce haploid spermatids, and (3) spermiogenesis, where round spermatids undergo morphological changes, such as nuclear condensation and elongation and flagella formation, to become spermatozoa [2, 3]. Spermatozoa released from seminiferous tubules are collected in the rete testis and transported to the caput epididymis via efferent ducts (EDs) (Fig 1A) [4, 5]. Spermatozoa are initially immotile and gradually mature and gain full motility as they travel through the epididymis, where they are stored in the cauda epididymis for release [6, 7].

**Fig 1.**
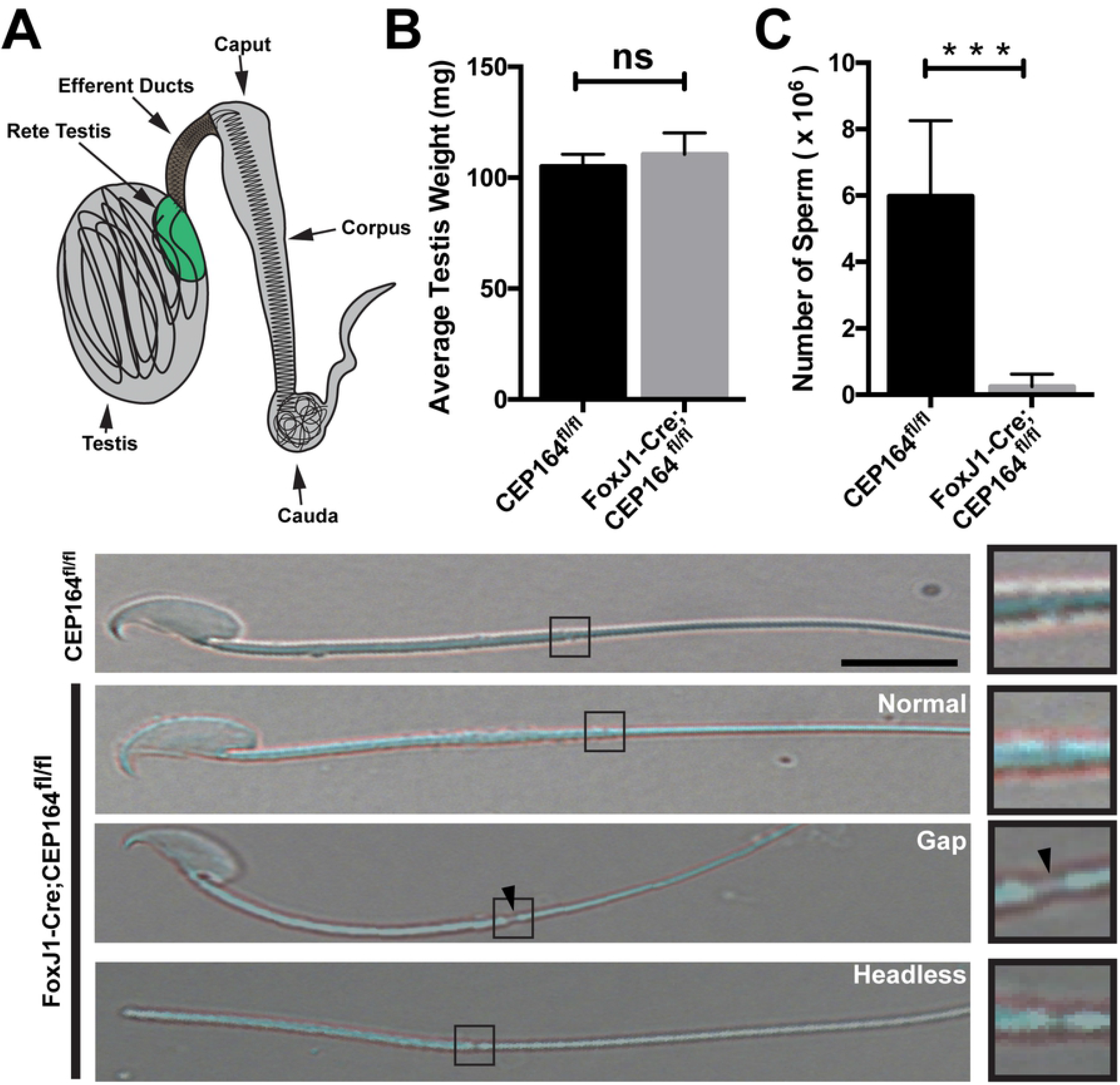
Loss of CEP164 leads to reduced sperm counts and aberrant sperm morphology. (A) Schematic of the male reproductive system. (B) Testes were isolated from 2 to 4-month-old CEP164^fl/fl^ and FoxJ1-Cre;CEP164^fl/fl^ mice and weighed (N=5 per genotype). Error bars represent means ± SD. ns, not significant. (C) Sperm were collected from the cauda epididymis of 2 to 4-month-old mice and counted on a hemocytometer (N=5 per genotype). Error bars represent means ± SD. ***, p<0.001. (D) Representative images of isolated sperm. The boxed regions are magnified on the right. Arrowheads point to a gap between the midpiece and the principal piece. Scale bar, 5 μm.

Sertoli cells are somatic epithelial cells that line the seminiferous tubule and play critical roles in spermatogenesis, production of seminiferous fluid, and phagocytosis of degenerated germ cells [8]. The seminiferous fluid secreted from Sertoli cells propels immotile sperm into the ED. In the EDs, nonciliated cells reabsorb up to 90% of the Sertoli cell secretions in order to concentrate the luminal content as sperm transit into the epididymis [9, 10]. Multiciliated cells (MCCs) stir the luminal fluid to prevent sperm agglutination and coagulation [4, 5]. Lastly, smooth muscle cells in the periphery of EDs assist in transporting immotile spermatozoa into the caput epididymis via peristaltic contractions [11]. Once in the epididymis, the spermatozoa continue to develop as they transit from the caput, through the corpus, and finally to the cauda epididymis where they are stored for release (Fig 1A).

Cilia are evolutionarily conserved, microtubule-based organelles that play crucial roles in a multitude of cellular processes [12–14]. As cells exit the cell cycle, cilia are assembled on the apical cell surface from the basal body, which is derived from the mother centriole [15]. Mother centrioles contain accessory structures, including subdistal and distal appendages at their distal end, whereas daughter centrioles do not. While subdistal appendages are involved in microtubule anchoring, distal appendages are important for the recruitment of small vesicles and subsequent docking of basal bodies to apical membranes [16–18]. Cilia are broadly categorized into two distinct groups: primary and multi-cilia. Primary cilia show a characteristic 9+0 microtubule arrangement and are present on the surface of many different cell types [19]. Primary cilia are involved in mechanosensation and signal transduction, most notably, Hedgehog signaling [13]. In contrast, multicilia, which have a 9+2 microtubule arrangement, typically move in a sweeping motion to help clear debris from airways, transport the ovum through the fallopian tube, and circulate cerebrospinal fluid [20–22]. As a notable exception, multicilia in EDs have recently been shown to produce a coordinated, whip-like motion, generating a strong rotational force in the lumen to prevent sperm agglutination [11]. The dysfunction of cilia is associated with a wide range of human genetic diseases, known as ciliopathies, including polycystic kidney disease, retinal degeneration, airway infection, and infertility [23–25].

Male infertility has been reported in mouse models with defective multicilia. The geminin family members, GEMC1 and MCIDAS, are essential for the MCC differentiation program [26–30]. GEMC1 and MCIDAS function as transcriptional activators in a complex with E2F4/5 and DP1 transcription factors to activate the expression of numerous ciliary genes including *FoxJ1* and *Cyclin O* (*Ccno*) [28, 31, 32]. It has been shown that male mice lacking GEMC1, MCIDAS, or CCNO are infertile, resulting from MCC defects in EDs [33]. Similarly, mice deficient for E2F4 and E2F5 or the microRNA clusters miR-34/449 exhibit loss of multicilia in EDs and male infertility [11, 34]. These mouse models display common pathological features including dilation of the seminiferous tubules and rete testis, sperm accumulation in EDs, and a lack of sperm in the epididymis. It is currently thought that MCCs in the ED epithelium stir the luminal contents to prevent sperm agglutination [11]. Without functional MCCs in EDs, sperm aggregate, ultimately leading to ED obstructions. This may cause fluid retention and back pressure in seminiferous tubules, resulting in testicular atrophy and infertility. In sum, these findings point to a crucial role for MCCs in male fertility.

CEP164, centrosomal protein of 164 kDa, localizes to distal appendages and plays an essential role in basal body docking and ciliogenesis [16–18, 35]. We demonstrated that CEP164 binds Chibby1 (Cby1) and facilitates the basal body recruitment of a protein complex of Cby1 and Cby1-interacting Bin/Amphiphysin/Rvs (BAR) domain-containing 1 and 2 (ciBAR1 and 2) (also known as FAM92A and B) [17, 36]. The Cby1-ciBAR complex then promotes the formation of membranous structures, called ciliary vesicles, which are critical for subsequent basal body docking to the apical cell membrane, during airway MCC differentiation. More recently, we reported that mice lacking CEP164 in MCCs (FoxJ1-Cre;CEP164^fl/fl^) show a significant loss of multicilia in the trachea, oviduct, and ependyma, and about 20% die due to profound hydrocephalus. In tracheal MCCs, the basal body recruitment of Cby1 and ciBAR proteins was severely diminished in the absence of CEP164 [17]. Furthermore, we found that FoxJ1-Cre;CEP164^fl/fl^ males are completely infertile. However, the precise role of CEP164 in the male reproductive system has remained unexplored.

Here, we report the characterization of male infertility in FoxJ1-Cre;CEP164^fl/fl^ mice. We found that, in these mice, the seminiferous tubules and rete testis show substantial dilation, reminiscent of dysfunctional multicilia in EDs. In addition, sperm accumulation and agglutination were commonly observed in the seminiferous tubules and EDs, eventually leading to a blockage of sperm progression and a dramatic reduction in the number of sperm in the cauda epididymis. Consistent with these findings, multicilia were barely detectable in the EDs of FoxJ1-Cre;CEP164^fl/fl^ mice although FoxJ1-positive immature cells were observed. Furthermore, transmission electron microscopy (TEM) studies revealed the presence of numerous undocked basal bodies in the cytoplasm of MCCs in the EDs of FoxJ1-Cre;CEP164^fl/fl^ mice. All together, our findings suggest that ablation of CEP164 in MCCs results in basal body docking defects and loss of multicilia in EDs, leading to male infertility.

## Results

### FoxJ1-Cre;CEP164^fl/fl^ mice show low sperm counts and abnormal sperm morphology

We have previously reported that FoxJ1-Cre-mediated deletion of CEP164 leads to fully penetrant male sterility [17]. To begin to understand the function of CEP164 in male fertility, we first examined testis weights and sperm counts. We found no significant difference in testis weights between control CEP164^fl/fl^ and FoxJ1-Cre;CEP164^fl/fl^ mice (Fig 1B). However, there was a dramatic decrease in the number of sperm from the cauda epididymis in FoxJ1-Cre;CEP164^fl/fl^ mice (Fig 1C), yet these sperm were motile. Interestingly, some of the sperm from FoxJ1-Cre;CEP164^fl/fl^ mice displayed morphological defects such as detached heads and gaps between the midpiece and the principal piece (Fig 1D).

### Aberrant morphology of the rete testis and cauda epididymis in FoxJ1-Cre;CEP164^fl/fl^ mice

Histological analyses of testes from FoxJ1-Cre;CEP164^fl/fl^ mice revealed marked luminal dilation of seminiferous tubules with thinning of the epithelium (Fig 2A). Elongated flagella were visible in the lumen of seminiferous tubules in FoxJ1-Cre;CEP164^fl/fl^ mice (bottom panels). In the control CEP164^fl/fl^ cauda epididymis, tubules were filled with spermatozoa (Fig 2B). In stark contrast, many cauda epididymal tubules in FoxJ1-Cre;CEP164^fl/fl^ mice contained only few sperm with abnormal fluid secretions (i and iii). Commonly, we also observed a few tubules containing agglutinated spermatozoa (asterisks and ii).

**Fig 2.**
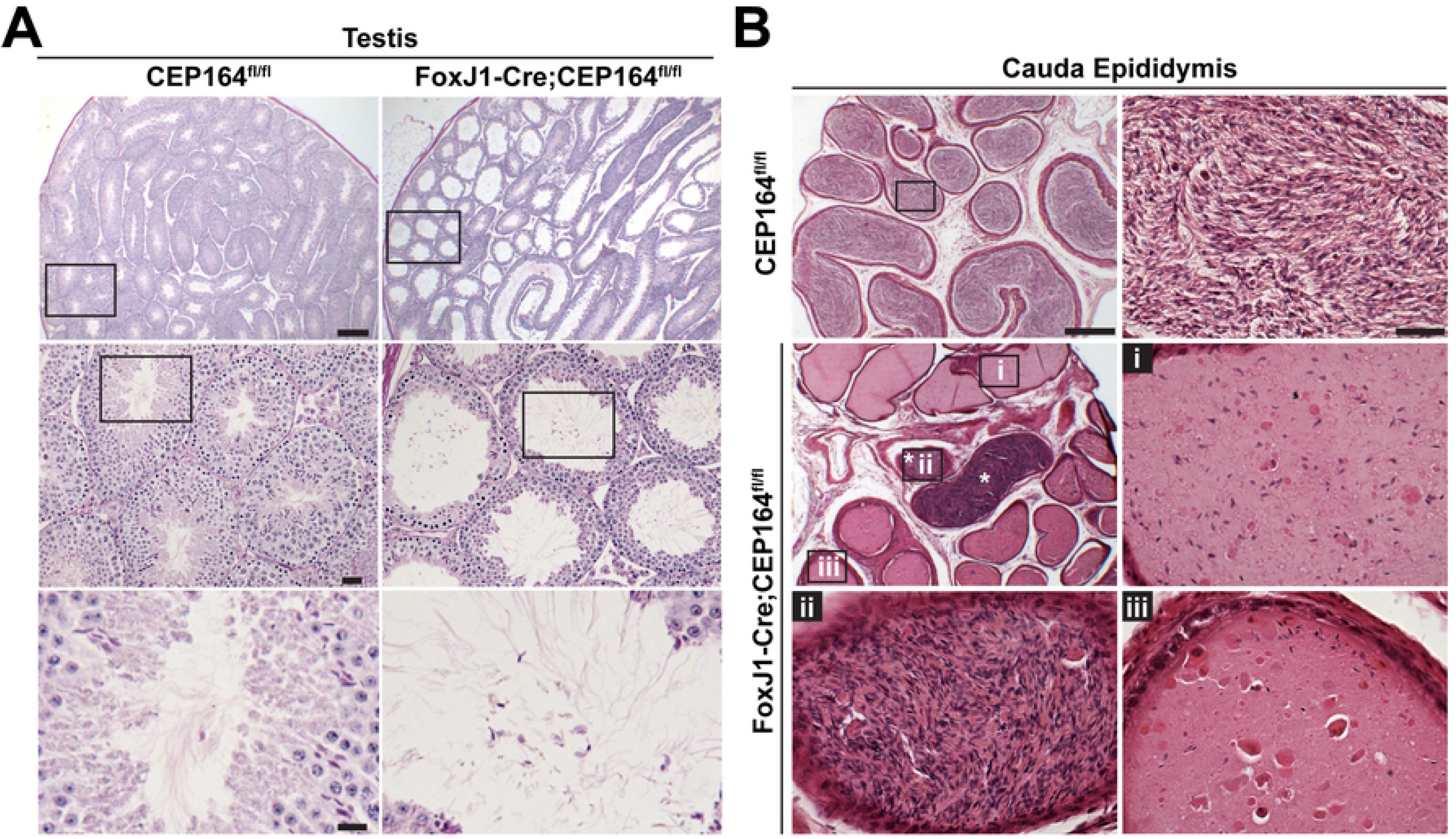
Deletion of CEP164 results in morphological changes in the male reproductive tract. (A) PAS staining of adult testicular sections. The boxed areas are enlarged in lower panels. Scale bars, 200 μm in the top, 25 μm in the middle, and 10 μm in the bottom panels. (B) H&E staining of cauda epididymal sections. The boxed areas are shown at high magnification. Scale bars, 200 μm and 25 μm (magnified images).

The thinning of the seminiferous epithelium is a hallmark of dysfunctional multicilia in the efferent ducts (EDs) [33]. It is typically associated with dilation of the rete testis, which connects seminiferous tubules to EDs (Fig 1A). Histological assessment of the rete testis revealed substantial dilation in FoxJ1-Cre;CEP164^fl/fl^ mice compared to that of control CEP164^fl/fl^ mice (Fig 3A, arrows). Intriguingly, clogged seminiferous tubules were frequently visible near the rete testis in intact testes from FoxJ1-Cre;CEP164^fl/fl^ mice (Fig 3B, arrowheads). Cross sections of the clogged seminiferous tubules confirmed extensive sperm agglutination (Fig 3C, arrowheads). Sperm agglutination was detected in all 6 mice that we examined. Taken together, FoxJ1-Cre;CEP164^fl/fl^ mice exhibit significant dilation of the seminiferous tubules and rete testis and sperm agglutination, most likely due to defective multicilia in EDs.

**Fig 3.**
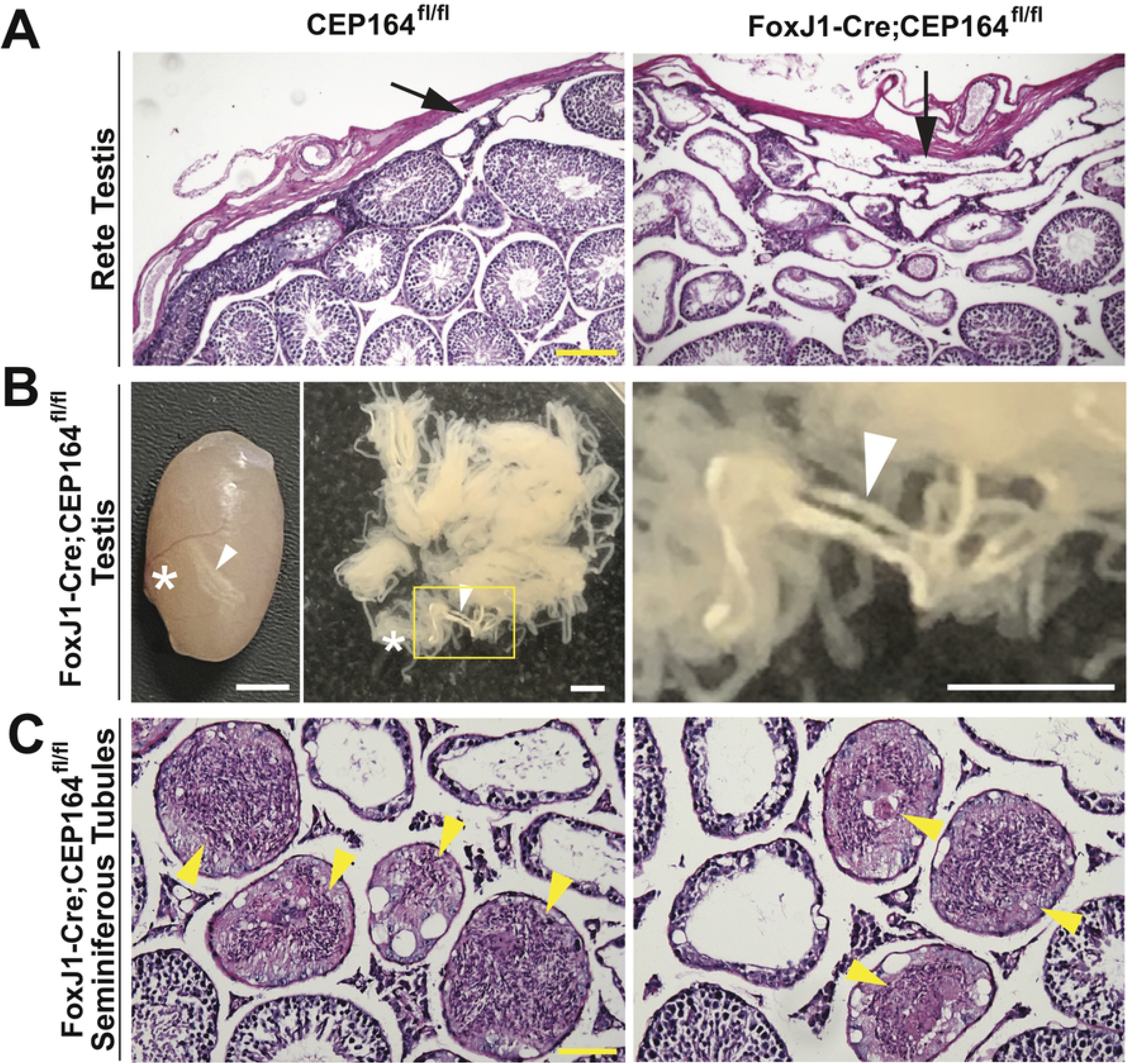
Rete testis dilation and sperm agglutinations are common in FoxJ1-Cre; CEP164^fl/fl^ mice. (A) PAS staining of the rete testis from adult mice. Arrows point to the rete testis. Scale bar, 100 μm. (B) Left: a representative testis from FoxJ1-Cre;CEP164^fl/fl^ mice, showing clogged seminiferous tubules. Right: dispersed seminiferous tubules. The boxed area is magnified on the right. Arrowheads point to clogged seminiferous tubules, and asterisks indicate the location of the rete testis. Scale bars, 2 mm. (C) PAS staining of testicular sections from FoxJ1-Cre;CEP164^fl/fl^ mice, showing sperm agglutination in seminiferous tubules (arrowheads). Scale bar, 50 μm.

### CEP164 localizes to the basal bodies of multiciliated cells in efferent ducts and sperm flagella in the testis

CEP164 localizes to the base of both primary and multi-cilia and plays essential roles in ciliogenesis in various cell types [17, 18, 35]. Using *CEP164-lacZ* reporter mice, we demonstrated that CEP164 is ubiquitously expressed in the mouse testis [17]. To investigate the subcellular distribution of CEP164 in the male reproductive tract, we performed immunofluorescence (IF) staining. The EDs join the testis to the epididymis, and sperm are concentrated as they progress from the rete testis into the caput epididymis (Fig 1A) [5, 9]. Multiciliated cells (MCCs) are present in the ED epithelium and have been shown to play a critical role in preventing the agglutination of sperm by stirring the luminal fluids [11]. In control CEP164^fl/fl^ mice, CEP164 localized to the apical surface of the ED epithelium (Fig 4A). No such apical CEP164 signals were detectable in FoxJ1-Cre;CEP164^fl/fl^ EDs, confirming efficient Cre-mediated recombination and antibody specificity. We also consistently noticed dilation of EDs to varying degrees. Intriguingly, costaining with the lectin peanut agglutinin (PNA), which binds to the outer acrosomal membrane of sperm [37], showed robust accumulation of sperm in the EDs of FoxJ1-Cre;CEP164^fl/fl^ mice, although no sperm were typically present in those of control CEP164^fl/fl^ mice due to rapid transport of sperm through EDs (Figs 4A and S1). Moreover, we found that CEP164 localized to the basal bodies of elongating flagella, marked by the basal body marker γ-tubulin (G-tub), in spermatids in both control CEP164^fl/fl^ and FoxJ1-Cre;CEP164^fl/fl^ testes (Fig. 4B), indicating that CEP164 is involved in flagella formation. It has been reported that FoxJ1 and FoxJ1-promoter-driven Cre are expressed in spermatids as early as round-spermatid stages (Fig S2) [38–40]. This raised the possibility that FoxJ1-Cre-mediated recombination at the *CEP164* locus is inefficient in spermatids. To examine this, we counted the number of CEP164-positive basal bodies in late-stage spermatids (steps 12 to 13). Quantification analysis revealed no significant differences in the percentage of CEP164-positive basal bodies between CEP164^fl/fl^ (89 ± 2%) and FoxJ1-Cre;CEP164^fl/fl^ (85 ± 4%) testes (Fig 4B).

**Fig 4.**
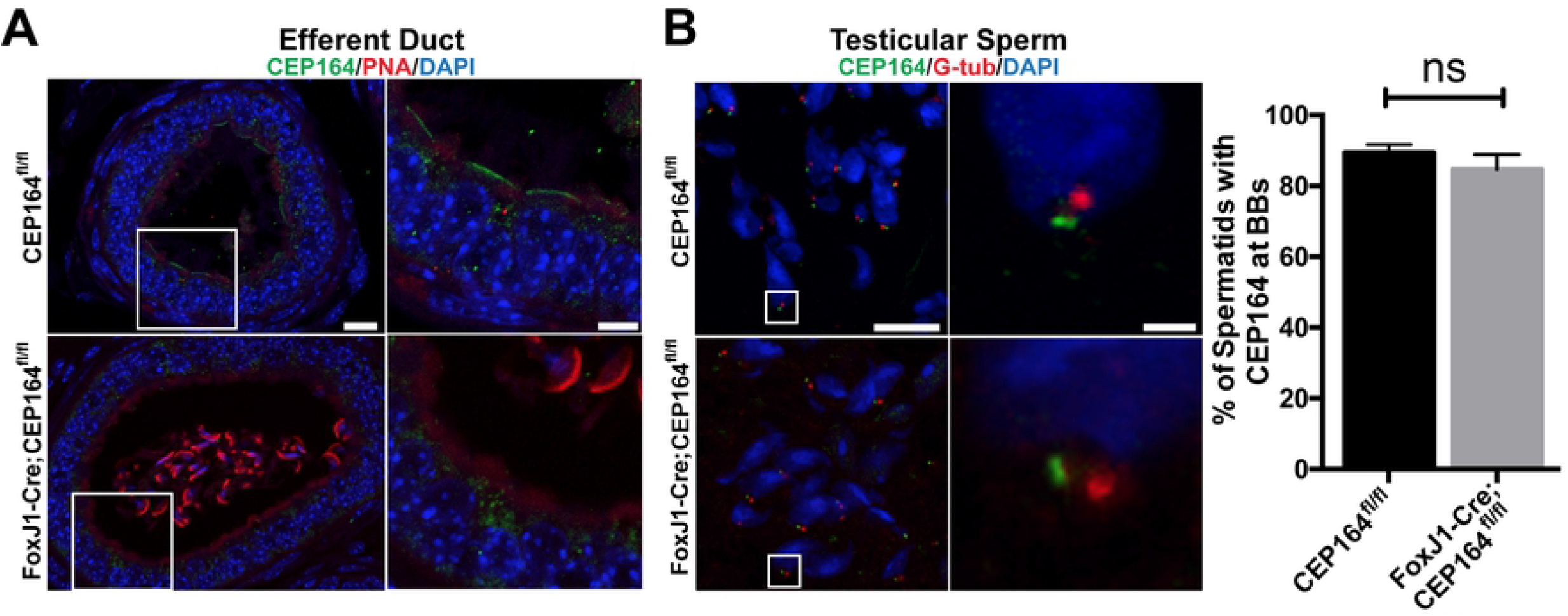
CEP164 expression in the male reproductive tract. (A) IF staining of ED sections for CEP164 and the lectin PNA. The boxed areas are magnified on the right. Scale bars, 10 μm and 5 μm (magnified images). Note that many sperm are typically found in the EDs of FoxJ1-Cre;CEP164 mice. (B) IF staining of testicular sections for CEP164 and the basal body marker G-tub. The boxed areas are magnified on the right. Scale bars, 10 μm and 1 μm (magnified images). Quantification of the percentage of spermatids with CEP164 at G-tub-positive basal bodies is shown on the right. A total of 400 basal bodies were counted in S12 to S13 spermatids for each genotype (N=3 seminiferous tubules). Error bars represent means ± SD. ns, not significant. Nuclei were detected with DAPI.

### Loss of muticilia in the efferent duct epithelium of FoxJ1-Cre;CEP164^fl/fl^ mice

Our data so far point to the possibility that defective multicilia in the EDs of FoxJ1-Cre;CEP164^fl/fl^ mice lead to sperm agglutination, profound dilation of the rete testis and seminiferous tubules, and ultimately infertility. To directly test this, we performed histological analyses of EDs. As expected, the EDs of control CEP164^fl/fl^ mice showed abundant multicilia with a length of 10 to 15 μm, extending into the lumen (Fig 5A). In contrast, multicilia were hardly detectable in the EDs of FoxJ1-Cre;CEP164^fl/fl^ mice. We also frequently observed the agglutination of sperm in the ED lumem of FoxJ1-Cre;CEP164^fl/fl^ mice, although control CEP164^fl/fl^ EDs typically contained no or only few sperm (Fig 5A). IF staining for the ciliary axoneme marker acetylated α-tubulin (A-tub) demonstrated a nearly a complete loss of multicilia in the EDs of FoxJ1-Cre;CEP164^fl/fl^ mice (Fig 5B), confirming the histological analysis.

**Fig 5.**
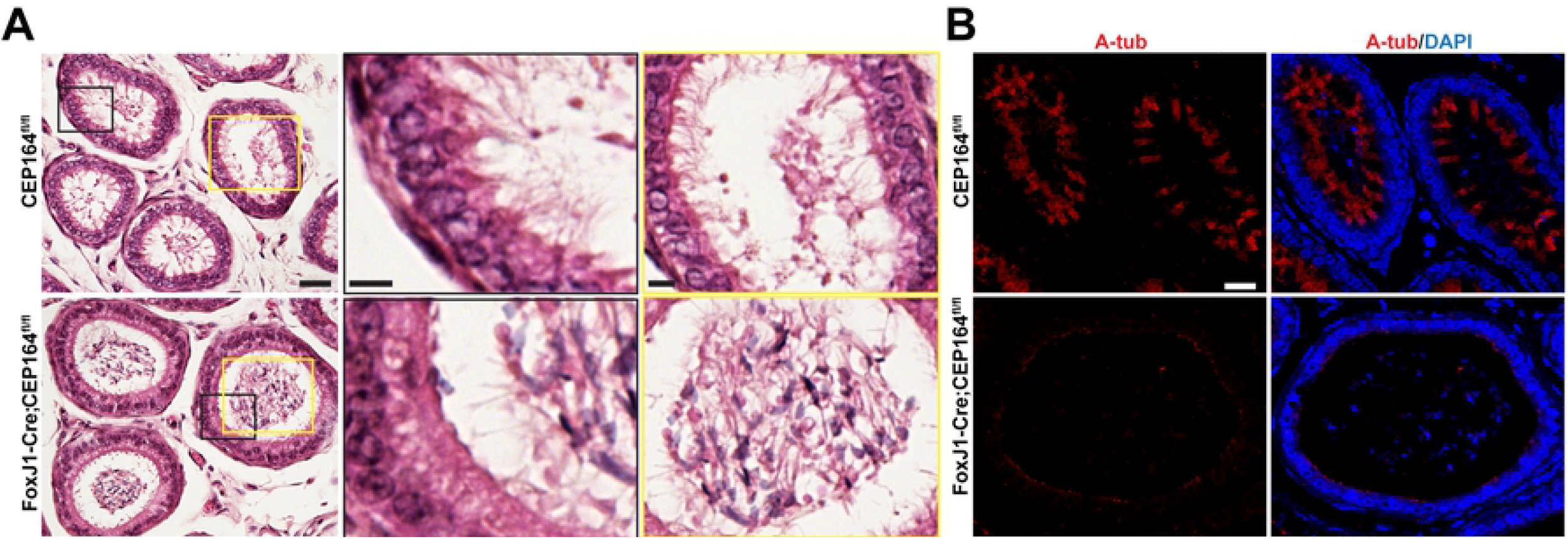
CEP164 is required for ciliogenesis in the efferent duct. A) H&E staining of ED sections. The boxed areas in black are magnified in the middle, and the boxed areas in yellow are magnified on the right. Scale bars, 20 μm and 5 μm (magnified images). (B) IF staining of ED sections for the ciliary axoneme marker A-tub. Nuclei were detected with DAPI. Scale bar, 20 μm.

To examine if ED progenitor cells are committed to the MCC lineage in the absence of CEP164, we performed IF staining for the MCC marker FoxJ1 [38, 39, 41]. Similar to control CEP164^fl/fl^ EDs, FoxJ1 was detected in the nuclei of the ED epithelium in FoxJ1-Cre;CEP164^fl/fl^ mice, but these cells lacked multicilia (Fig 6A and 6B). Quantification of FoxJ1-positive MCCs in FoxJ1-Cre;CEP164^fl/fl^ EDs revealed that 98.8% of MCCs were devoid of multicilia (494 out of 500 MCCs), indicating a high efficiency of Cre-mediated recombination. The remaining A-tub-positive structures on the apical cell surface were reminiscent of primary cilia of secretory cells (Fig 6B, arrowheads) [5, 42]. These primary cilia are always located near the tight junctional complex at the apical cell surface [5]. To verify this, we performed IF staining for A-tub and the secretory cell marker aquaporin 1 (AQP1). In control CEP164^fl/fl^ EDs, IF staining for AQP1 showed apical enrichment in secretory cells (Fig. 6C, arrow). These cells were clearly present in the EDs of FoxJ1-Cre;CEP164^fl/fl^ mice, and, as predicted, the A-tub-positive structures coincided with AQP-1-positive secretory cells (Fig 6C, arrowheads).

**Fig 6.**
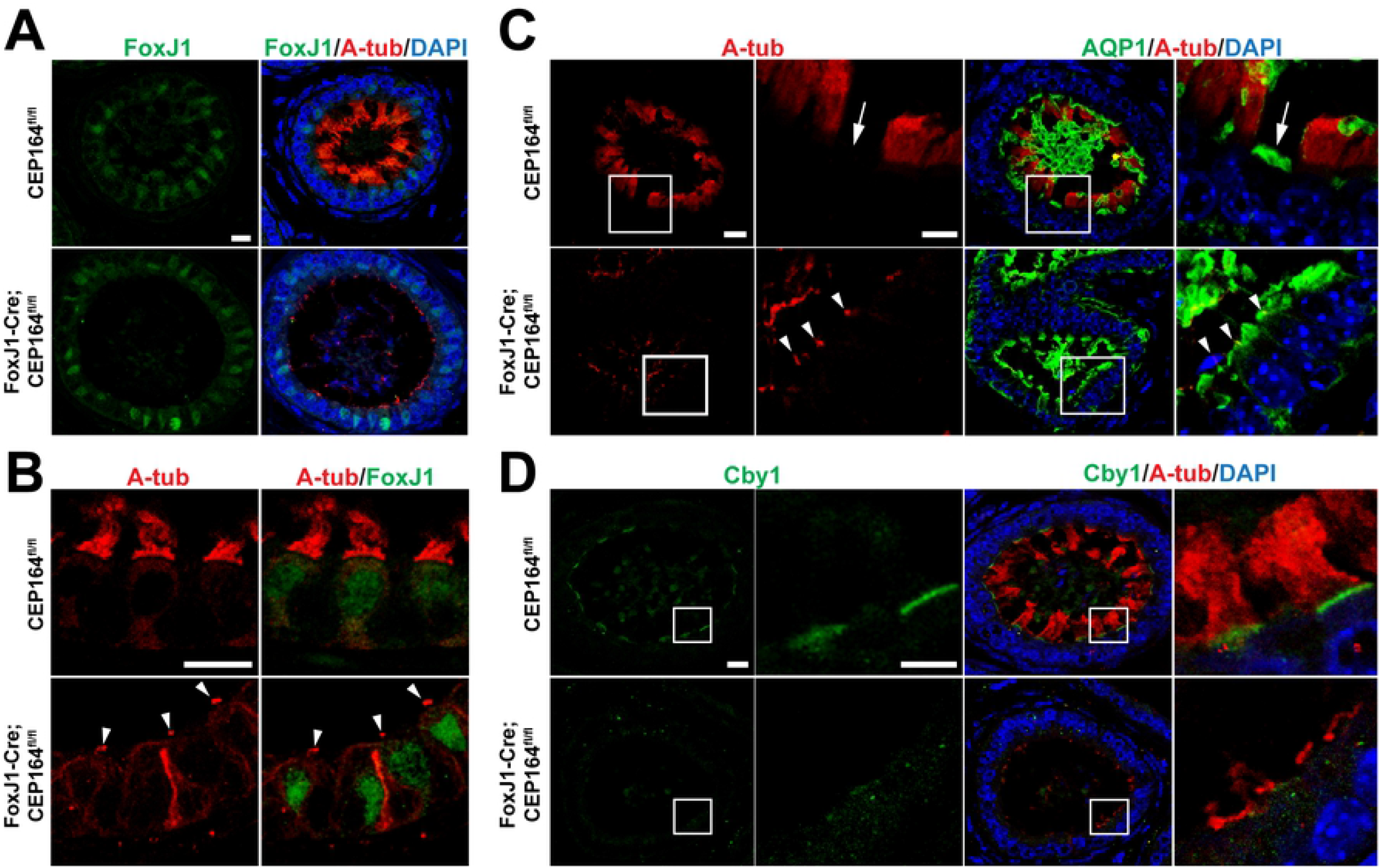
Marker analysis of the efferent ductal epithelium. (A) IF staining of ED sections for A-tub and the MCC marker FoxJ1. Scale bar, 10 μm. (B) IF staining of ED sections for A-tub and FoxJ1. Arrowheads point to the primary cilia of secretory cells in FoxJ1-Cre;CEP164 EDs. Scale bar, 10 μm. (C) IF staining of ED sections for A-tub and the secretory cell marker aquaporin 1 (AQP1). The boxed areas are magnified on the right. Arrows denote AQP1-positive secretory cells. Arrowheads point to the primary cilia of secretory cells. Scale bars, 10 μm and 5 μm (magnified images). (D) IF staining of ED sections for Cby1, a CEP164-binding basal body protein, and A-tub. Note that Cby1 is lost from the apical surface of MCCs in FoxJ1-Cre;CEP164 efferent ducts. The boxed areas are enlarged on the right. Scale bars, 10 μm and 5 μm (enlarged images). Nuclei were detected with DAPI.

Previously, we demonstrated that CEP164 physically interacts with and recruits Cby1 to the distal appendages of nascent basal bodies to facilitate the efficient formation of ciliary vesicles during the early stages of airway MCC differentiation, and both remain at the base of mature cilia [16, 17, 43]. We observed Cby1 at the apical surface of MCCs, basal to A-tub staining, in control CEP164^fl/fl^ EDs. However, apical Cby1 staining was lost in FoxJ1-Cre;CEP164^fl/fl^ EDs (Fig 6D). Similarly, the apical localization of NPHP1, a maker for the ciliary transition zone that acts as a diffusion barrier at the ciliary base [44, 45], was also altered in FoxJ1-Cre;CEP164^fl/fl^ EDs (Fig S3). These results are consistent with basal body docking defects.

### CEP164 is critical for basal body docking in multiciliated cells in efferent ducts

CEP164 plays pivotal roles in recruitment of small preciliary vesicles to distal appendages to assemble ciliary vesicles, thereby promoting basal body docking [16–18]. In airway MCCs, loss of CEP164 is associated with defects in basal body docking and subsequent cilium elongation [17]. To investigate if this is also the case for MCCs in the EDs of FoxJ1-Cre;CEP164^fl/fl^ mice, we performed transmission electron microscopy (TEM) analyses. In control CEP164^fl/fl^ EDs, we observed basal bodies docked to the apical cell surface with multicilia extending into the lumen (Fig 7, arrowheads). In FoxJ1-Cre;CEP164^fl/fl^ EDs, however, many basal bodies were found undocked deep into the cytoplasm (Fig 7, arrowheads). These results underscore the key role of CEP164 in basal body docking.

**Fig 7.**
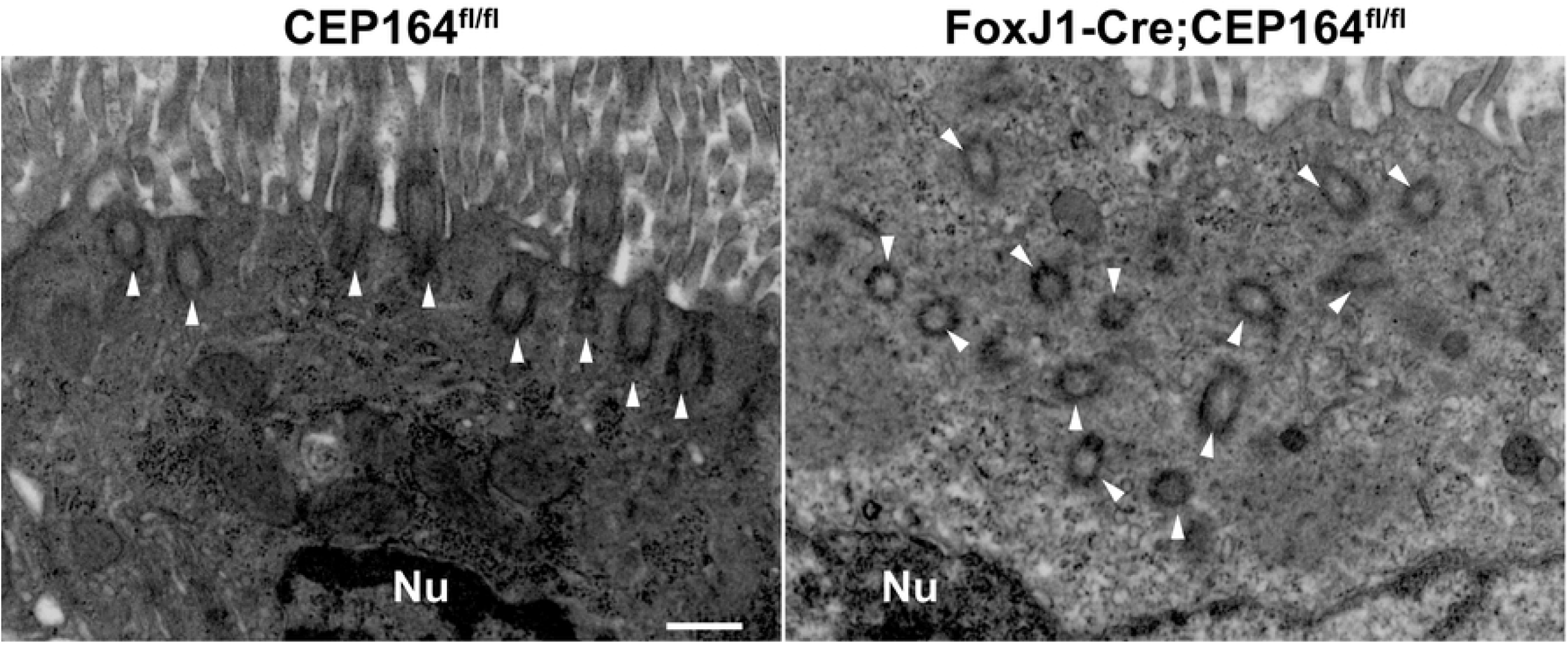
CEP164 is necessary for basal body docking in multiciliated cells in efferent ducts. Shown are representative TEM images of MCCs in EDs. Arrowheads point to basal bodies. All 29 MCCs examined from the EDs of FoxJ1-Cre;CEP164 mice displayed severe basal body docking defects, whereas all 31 MCCs in those of control CEP164^fl/fl^ mice showed basal bodies properly docked at the apical surface. Nu, nucleus. Scale bar, 500 nm.

## Discussion

MCCs are found in a restricted set of tissues, such as the airway, oviduct, spinal cord, and brain ventricles and play crucial roles in fluid propulsion across the epithelial surface. A growing body of evidence suggests that MCCs in EDs play a major role in male fertility [11, 33, 34]. It has recently been shown that these multicilia act as agitators that generate strong turbulence to maintain immotile spermatozoa in suspension in the ED lumen [11]. Upon loss of these multicilia, aggregation of sperm leads to ED occlusions, thereby preventing the transport of sperm into the epididymis.

We found that loss of CEP164 in MCCs leads to male infertility due to a nearly complete ablation of multicilia in the ED epithelium. Consequently, FoxJ1-Cre;CEP164^fl/fl^ mice consistently develop sperm agglutinations in the seminiferous tubules, EDs, and cauda epididymis (Figs 2, 3, 5, and S1). In particular, large stretches of seminiferous tubules near the rete testis were visibly clogged (Fig 2B). Ultimately, this results in dilation of the rete testis and seminiferous tubules, followed by testicular degeneration. Thus, it indicates that dysfunctional multicilia in EDs leads to sperm agglutinations and obstruction of sperm passage in different segments of the male reproductive tract.

The impact of CEP164 deletion on MCC differentiation in EDs is prominent as multicilia are barely detectable (Figs 5–7). We demonstrated that the recruitment of basal body proteins to the apical surface is abolished, and many basal bodies are undocked in the EDs of FoxJ1-Cre;CEP164^fl/fl^ mice. Our findings therefore confirm the key role of CEP164 in basal body docking during MCC differentiation *in vivo*.

A recent single-cell RNA sequencing analysis suggests that CEP164 is ubiquitously expressed throughout mouse spermatogenesis (Fig S2) [40]. Our IF staining demonstrated that CEP164 intensely localizes to the base of elongating flagella in spermatids (Fig 4B), suggesting that CEP164 is involved in flagellogenesis. This is in contrast to the fly CEP164 homolog (CG9170), which is not expressed in the testis (Flybase). A recent study has shown that CEP164 also localizes to the capitulum and striated columns of the human sperm neck [46], although its functional significance is unknown. Thus, it is likely that CEP164 plays multiple roles during spermatogenesis.

Although CEP164 is still present at the base of sperm flagella in FoxJ1-Cre;CEP164^fl/fl^ mice at comparative levels to control (Fig 4B), some spermatozoa collected from the cauda epididymis show morphological defects such as detached heads and a gap between the midpiece and principal piece of the sperm tail (Fig 1D). One possible explanation is that fluid retention and back pressure, caused by ED obstructions, may contribute to the sperm morphological defects. Alternatively, it could be attributable to changes in the luminal microenvironment of the epididymis as testicular sperm undergo complex maturation processes as they travel through the epididymis [47–49].

In summary, results presented here provide compelling evidence that CEP164 is necessary for proper MCC differentiation in EDs and male fertility. CEP164 localizes to the base of flagella, suggesting that CEP164 also plays a role in flagellogenesis. The roles of CEP164 in the testis are unclear and awaits further investigation using tissue and cell type-specific Cre driver lines. Primary ciliary dyskinesia (PCD) is the most prominent ciliopathy associated with the dysfunction of multicilia [50, 51]. Male infertility and sub-fertility are common manifestations of PCD. In addition to sperm flagellar defects, compromised functions of multicilia in EDs may, at least in part, contribute to the male infertility of PCD patients. Our FoxJ1-Cre;CEP164^fl/fl^ mouse model provide a powerful tool to further dissect the role of multicilia in the male reproductive system.

## Methods and Materials

### Ethics statement

All mice were handled in accordance with NIH guidelines, and all protocols were approved by the Institutional Animal Care and Use Committee (IACUC) of Stony Brook University (#2010–1393).

### Generation of FoxJ1-Cre;CEP164^fl/fl^ mice

FoxJ1-Cre;CEP164^fl/fl^ mice were generated as previously described [17]. Adult male mice at 2 to 4 months of age were used throughout the study. Genotyping was performed on tail biopsies using PCR primers: CEP164 KO-first, 5’-CCATCTGTCCAGTACCATTAAAAA-3’ and 5’-CCCAGAATACAACATGGGAGA-3’ (WT allele, 215 bp; floxed allele, 415 bp); Cre, 5’-CGTATAGCCGAAATTGCCAGG-3’and 5’-CTGACCAGAGTCATCCTTAGC-3’ (327 bp).

### Sperm counts and imaging

One cauda epididymis was dissected from 2-to 4-month-old mice and placed in a 35 mm dish with 500 μl PBS, pH 7.4. The tissue was then minced into small pieces and placed into a 37°C incubator for 30 minutes. The sperm were collected, and the dish was washed with 500 μl of PBS. The sperm were pooled and counted on a hemocytometer with appropriate dilutions. In addition, the sperm were centrifuged and fixed in 4% paraformaldehyde (PFA) overnight and mounted onto glass slides for image acquisition.

### Histological staining

Testes, EDs, and epididymides were dissected from adult mice and fixed in 4% PFA. The tissues were then embedded in paraffin blocks, and 5-μm sections were cut and stained with either hematoxylin and eosin (H&E) (Poly Scientific R&D Corp) or periodic acid-Schiff reagent (PAS) (Sigma-Aldrich) and mounted with Permount (Fischer Scientific).

### Immunofluorescence staining

For IF staining of PFA-fixed paraffin embedded tissues, tissue sections were deparaffinized and rehydrated, and then antigen retrieval was performed using sodium citrate buffer (10 mM sodium citrate, 0.05% Tween 20, pH 6.0) in an autoclave for 17.5 minutes. The specimens were then subjected to extraction using 0.5% Triton X-100 in PBS for 5 minutes, followed by blocking for 1 hour at room temperature using 5% goat serum in antibody dilution solution [2% bovine serum albumin (BSA) in PBS]. Primary antibodies were applied overnight at 4°C, and then secondary antibodies were applied for 1 hour at room temperature in antibody dilution solution. Finally, DAPI was applied for 3 minutes at room temperature, and the specimens were mounted with Fluoromount-G (Southern Biotech). For primary antibodies used, see supplemental table S1. The following secondary antibodies were used at a 1:200 dilution: goat anti-rabbit IgG conjugated with either DyLight 488 or DyLight 549 and horse anti-mouse IgG conjugated with either DyLight 488 or DyLight 549 (Vector Laboratories). The lectin PNA labeled with fluorescein or rhodamine was obtained from Vector Laboratories.

For IF staining for CEP164 and γ-tubulin at the base of flagella, freshly isolated testes were snap-frozen using 2-methylbutane and liquid nitrogen, and 10-μm thick sections were cut. The tissue was allowed to air dry for 30 minutes and then fixed with 4% PFA at room temperature, followed by acetone at −20°C for 10 minutes each. Subsequent steps were performed as described above.

### Generation and purification of NPHP1 antibody

A rabbit anti-NPHP1 antibody was generated at Open Biosystems, Inc against His-tagged human NPHP1 N-terminal region (aa1-153) containing coiled-coil and SH3 domains. The antigen was expressed and purified from *E. coli*. The antibody was purified by antigen affinity chromatography using a GST-NPHP1 (aa1-153) fusion protein and dialyzed against PBS.

### Image acquisition and analysis

For histological staining and sperm, images were acquired on a Leica DMI6000B microscope with either a 20X/0.50 NA, 40X/0.75 NA, or 63X/1.25 NA (oil) objective equipped with a DFC7000T camera. The sperm images were taken in differential interference contrast (DIC). For IF staining, confocal images were taken on a Leica SP8X with a HCPL APO 100X/1.4 NA oil objective. All images were analyzed using the Leica Application Suite X software, and final images were generated using ImageJ, Adobe Photoshop, and Adobe Illustrator.

### Transmission electron microscopy of efferent ducts

Samples used for TEM were processed using standard techniques as described previously [16, 17]. Briefly, adult EDs were fixed by immersion in 2% PFA and 2% glutaraldehyde in PBS overnight at 4°C. After fixation, samples were washed in PBS, placed in 2% osmium tetroxide in PBS, dehydrated in a graded series of ethanol, and embedded in Embed 812 resin (Electron Microscopy Sciences). Ultrathin sections of 80 nm were cut with a Leica EM UC7 ultramicrotome and placed on Formvar-coated slot copper grids. Sections were then counterstained with uranyl acetate and lead citrate and viewed with a FEI Tecnai12 BioTwinG^2^ electron microscope. Digital images were acquired with an XR-60 CCD digital camera system (Advanced Microscopy Techniques).

### Statistical Analysis

Two-tailed Student’s t-tests were used to determine statistical significance, and p < 0.05 was considered to be significant. Asterisks were used to indicate p values as follows: *** p < 0.001; and ns, not significant.

## Acknowledgement

We would like to acknowledge the Research Histology Core Laboratory in the Department of Pathology at Stony Brook University School of Medicine for assistance with histological preparations and Susan van Horn in the Central Microscopy Imaging Center Core at Stony Brook University for assistance with TEM.

---

**Fig S1. Sperm agglutination in the efferent ducts of FoxJ1-Cre; CEP164^fl/fl^ mice.**

IF staining of ED sections with the lectin PNA (green). Nuclei were detected by DAPI. The boxed areas are magnified on the bottom. Scale bars, 500 μm and 50 μm (magnified images). Note that profound sperm agglutinations are commonly found in the EDs of FoxJ1-Cre;CEP164 mice (arrowheads). Arrows point towards the testis (T) or the epididymis (E).

**Fig S2. RNA levels of FoxJ1 and CEP164 during spermatogenesis.**

The datasets for spermatogenic gene expression were retrieved from Ernst *et al*. [40] through their website (https://marionilab.cruk.cam.ac.uk/SpermatoShiny/). The data are represented by a standard box-whisker plot with black dots representing outliers. SG, spermatogonia; eP, early-pachytene spermatocyte; mP, mid-pachytene spermatocyte; lP, late-pachytene spermatocyte; D, diplotene spermatocytes; M, meiosis; and S1-11, step 1-11 spermatids.

**Fig S3. The ciliary transition zone marker NPHP1 is lost from the apical surface of efferent duct multiciliated cells in the absence of CEP164.**

IF staining of ED sections for A-tub and NPHP1. The boxed areas are magnified on the right. Scale bars, 10 μm and 5 μm (magnified images).

